# Generative adversarial networks in cell microscopy for image augmentation. A systematic review

**DOI:** 10.1101/2023.08.25.554841

**Authors:** Duway Nicolas Lesmes-Leon, Andreas Dengel, Sheraz Ahmed

## Abstract

Cell microscopy is the main tool that allows researchers to study microorganisms and plays a key role in observing and understanding the morphology, interactions, and development of microorganisms. However, there exist limitations in both the techniques and the samples that impair the amount of available data to study. Generative adversarial networks (GANs) are a deep learning alternative to alleviate the data availability limitation by generating nonexistent samples that resemble the probability distribution of the real data. The aim of this systematic review is to find trends, common practices, popular datasets, and analyze the impact of GANs in image augmentation of cell microscopy images. We used ScienceDirect, IEEE Xplore, PubMed, bioRxiv, and arXiv to select English research articles that employed GANs to generate any kind of cell microscopy images independently of the main objective of the study. We conducted the data collection using 15 selected features from each study, which allowed us to analyze the results from different perspectives using tables and histograms. 32 studies met the legibility criteria, where 18 had image augmentation as the main task. Moreover, we retrieved 21 publicly available datasets. The results showed a lack of consensus with performance metrics, baselines, and datasets. Additionally, we evidenced the relevance of popular architectures such as StyleGAN and losses including Vanilla and Wasserstein adversarial loss. This systematic review presents the most popular configurations to perform image augmentation. It also highlights the importance of design good practices and gold standards to guarantee comparability and reproducibility. This review implemented the ROBIS tool to assess the risk of bias, and it was not registered in PROSPERO.

## Introduction

The study of microorganisms is essential for progress in medicine, understanding them allows the development of drugs, treatments, and diagnosis of diseases, allowing mankind to improve not only their quality of life, but also their relationship with the environment [1]. Microorganisms cannot be seen with the naked eye, which had been a major limitation in the past. Fortunately, thanks to cell microscopy, researchers now count with several available tools that have relieved this limitation considerably.

Cell microscopy imaging is the set of techniques used to visualize microorganisms, it allows researchers to study anatomy, dynamics, and interactions between microorganisms. There are different microscopy techniques that have been developed to highlight specific features, some of the most popular techniques are optical, fluorescence and electron microscopy. However, each technique has intrinsic limitations, some have limited magnification, some require killing and fixing the samples, others require very specialized tools, making them inaccessible. Moreover, The preparation of the data carries its own challenges, including expensive and long sample preparation, environmental conditions that are not always reproducible, a limited time-window to visualize important events, ethical considerations depending on the source of the data, and time-consuming and prone-error analysis carried out by experts. The combination of all these factors make the cell imaging a domain with a significant impairment in the production of both raw and annotated data.

In the last years, deep learning (DL) has provided the field of computer vision with several alternatives to alleviate the challenges mentioned earlier. Training DL models in different tasks has several benefits that include decreasing image evaluation times, capturing information that is not easily seen to the naked eye, releasing specialized personnel from mechanical tasks, and simplifying their complexity. It is well known that model performance is proportional to the amount of data available to train; normally a significant amount of training data is required to achieve the best model generalization possible, and even though data is not a limitation for most fields, specialized domains such as cell microscopy struggle in this regard [2].

Fortunately, generative modeling is an area of machine learning whose task is to understand the distribution of data to produce synthetic samples resembling the real data distribution. Generative adversarial networks (GANs), a well established generative model and the focus of this study, comprise a family of DL algorithms based on the intuition of training two neural networks (NNs) simultaneously in a competitive fashion [3]. In computer vision, the evolution of GANs allowed researchers to propose promising solutions to several tasks comprising image classification, segmentation, augmentation, enhancements, and domain translation [4]. There are reviews analyzing GANs, their general foundation, and applications [4] [5], and specialized reviews of GANs in medical imaging [6] [7] [8] [9] [10]. Although some of these reviews compile some microscopy image datasets, we could not find a review focused on cell microscopy generation.

Considering the lack of data availability, we present a systematic review of GANs using cell microscopy imaging for image augmentation. The objectives of this work are:

- Analyze the studies using GANs to perform cell microscopy image augmentation.
- Analyze popular public available datasets to train generative models for cell microscopy image augmentation.
- Identify the most popular architectures, losses, and methods when using GANs in the field.
- Identify common practices of experimental design related to image augmentation.

We first screened different databases and selected relevant publications based on legibility criteria to analyze and summarize them to discover the main trends, challenges, and limitations present in the field. Moreover, we analyzed the publicly available datasets used in the selected publications and summarized their principal characteristics. Finally, we also briefly discuss the most representative GAN architectures and definitions for those readers with a biological background or enthusiastic researchers that are introducing themselves to the field of GANs. The main contributions of this work are:

- Detailed compilation of 32 selected studies from 5 different datasets, using features to facilitate their comparison.
- List and detailed description of the publicly available datasets used in the studies considered in this review.
- Important notions and considerations in the experimental design of generative modeling studies that are essential to produce research with robust methodology and results.

## Materials and methods

Before defining the guidelines for the systematic review, we included a brief section to present and discuss the basic notions of GANs for those readers that are not so familiar with them. We also present some representative losses and architectures that will be relevant in this study.

### Generative Adversarial Network (GAN)

The goal of GANs is to learn the probability distribution of a training dataset and use such distribution to generate new synthetic data that cannot be differentiated from the original dataset [11]. A GAN is traditionally composed of a generator that tries to learn the training data distribution, while the discriminator has to differentiate between synthetic and authentic data [3]. The training procedure of GANs can be conceived as a competition where the generator tries to fool the discriminator with realistic samples while the discriminator is trained to detect the generated ones. The formal definition of this problem is described as follows: with input noise *z ∈ Z* following the probability distribution *p_z_*and training data *x ∈ X* with probability distribution *p_data_*, two differentiable functions (generator and discriminator) represented as artificial neural networks (ANNs) are trained simultaneously.

The generator *G* takes *z* as input and transforms it into *x^i^ ∈ X^i^*that follows the generated probability distribution *p_g_*, the discriminator *D* takes input samples from both *X* and *X^i^* and returns a scalar representing the probability that its input belongs to *X*, if *D* = 0, then the discriminator concludes that such input does not belong to *X* and is, therefore, a synthetic sample.

The goal of *G* is to approximate *p_g_*to *p_data_*as much as possible using the information that *D* retrieves. Thus, it will be harder for *D* to find the synthetic samples once *G* is well-trained.

GANs have had a high development since first introduced by Goodfellow et al. in 2014 [3], several loss variants, architectures, and even additional components have been proposed to boost the performance of GANs. The Vanilla GAN loss implemented the cross-entropy loss, which has interesting properties that are discussed later. The objective is presented in the first row of Table 1.

**Table 1.** Summary of popular adversarial losses.

| Loss | Equation | Approximates to | Attributes |
| --- | --- | --- | --- |
| Vanilla | $\min_G \max_D \mathcal{L}(G, D) = \mathbb{E}_{x \sim p_{data}} [\log D(x)] + \mathbb{E}_{z \sim p_z} [\log(1 - D(G(z)))]$ | Jensen Shannon Divergence ($\mathcal{D}_{JS}$) with optimal discriminator | Robust against instability due to gradient exploit.<br>Instability due to gradient vanishing in case of disjoint supports. |
| Least-Square | $\min_D \mathcal{L}_{LS}(D) = \frac{1}{2} \mathbb{E}_{x \sim p_{data}} [(D(x) - b)^2]$<br>$+ \frac{1}{2} \mathbb{E}_{z \sim p_z} [(D(G(z)) - a)^2]$<br>$\min_G \mathcal{L}_{LS}(G) = \frac{1}{2} \mathbb{E}_{z \sim p_z} [(D(G(z)) - c)^2]$ | Pearson $\chi^2$ divergence ($\mathcal{D}_{\chi^2}$) with optimal discriminator | The least-square objective encourages $G$ to push the generated samples to the decision boundary of $D$.<br>Instability due to gradient vanishing in case of disjoint supports. |
| Wasserstein | $W(p_1, p_2) = \sup_{\vert \vert f\vert \vert _L \leq 1} \mathbb{E}_{x \sim p_1} [f(x)] - \mathbb{E}_{x \sim p_2} [f(x)]$ | Wasserstein-1 distance with K-Lipschitz | This objective measures the distance between distributions, making it robust against disjoint supports.<br>The discriminator is renamed as critic $C$ since it will no longer perform classification, $C$ outputs now a distance.<br>The critic output is correlated with image quality, a property that allows to monitor the training progress. |
| Relativistic | $\mathcal{L}_{rel}^D(C) = \mathbb{E}_{(x,z) \sim (p_{data}, Z)} [f_1(C(x) - C(G(z)))]$<br>$\mathcal{L}_{rel}^G(C) = \mathbb{E}_{(x,z) \sim (p_{data}, Z)} [f_1(C(G(z)) - C(x))]$ | N/A | This objective pretends to measure how much realistic is a generated sample compared to a real one.<br>Any discriminator performing classification can be transformed into a relativistic counterpart.<br>Considers prior knowledge related to the data and the training scheme that other losses do not consider. |

The Vanilla loss can be interpreted as the sum of the expected value of correctly classifying the real training samples and the expected value of correctly classifying the generated samples. Ideally, an optimal *G* causes *D* to output an average value of 0.5 no matter the data source (50% chance of being a real sample).

Additionally, Mirza and Osindero [12] extended the capacity of GANs with conditional GAN (cGAN). The idea of cGAN is simple yet powerful. It uses additional information *h* that encourages *D* to produce synthetic data containing the conditional information *h*. The benefit of cGAN is that *h* can be any data type used as additional input in both *G* and *D* as seen in Eq (1). One of the most popular conditional settings is to use an image as generator input. In this way, the GAN performs Image-to-Image (I2I) translation, a process where *G* pretends to transform images from an input domain into a target domain while preserving the meaning of the input image, e.g., transforming photographs into sketches. The most iconic architectures are Pix2Pix [13] and CycleGAN [14].

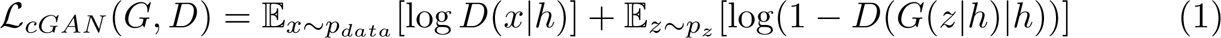

Later in time, new adversarial losses have been proposed with the goal of optimizing the limitations of Vanilla GAN, the most representative adversarial losses are Least-square GAN (LSGAN) [15] in 2017, Wasserstein GAN (WGAN) [16] in 2017, and Relativistic GAN (RGAN) [17] in 2019. Table 1 summarizes the main characteristics of these losses. For those readers interested in the formal definitions, training dynamics and theoretical implications of GANs, we encourage them to visit Arjovsky and Bottou’s work [18] in which they discussed all these topics thoroughly.

### StyleGAN

StyleGAN [19] is one of the most popular GAN architectures for image augmentation. The authors proposed a unique generator architecture inspired by style-transfer learning to better understand and control the synthesis process from the latent space where the generator samples its inputs. *G* is composed of two networks; a mapping, and a synthesis network. The mapping network is an 8-layer multilayer perception (MLP) that embeds the random noise input *z ∈ Z* into an intermediate latent space denoted as *W*. Different from other generators, the synthesis network’s input is a known constant and each convolutional block is fed with broadcasted noise to provide stochasticity and styles to control its adaptive instance normalization (AdaIN).

Based on the hypothesis that producing realistic outputs is easier from a disentangled space, the generator encourages the mapping network to produce a disentangled latent space *W* with more linear subspaces that facilitate the image synthesis. *w ∈ W* passes through learned affine transformations to produce the styles, Then, each style is fed into a specific block in the synthesis network. The empirical results indicated that, thanks to AdaIN, styles control the global features in the final output. Moreover, styles that are fed early in the network control high-level features, while styles fed deeper in the network define the finer features. On the other hand, the role of noise is to introduce stochastic variation that every image must have (such as hair disposition or freckles in human faces) while preserving the overall composition and high-level features produced by the styles. Later, StyleGAN had updates with StyleGAN2 [20] and StyleGAN3 [21]; different aspects of the architecture are revisited to correct artifacts and improve the image quality. StyleGAN and its updates achieved state-of-the-art results in human faces generation when each was published, making it one of the gold standard architectures in generative modeling.

### PathologyGAN

Quiros, Murray-Smith, and Yuan [22] used GANs to better understand the phenotypic variability of histological images. They part from the assumption that a GAN capable of producing realistic images should learn a disentangled latent space able to describe phenotypic features that are not easily seen. The authors used BigGAN [23] as the baseline of PathologyGAN and from there, they included features of StyleGAN including the mapping network, AdaIN, and style mixing to enforce a disentangled latent space. Finally, the architecture is trained using average relativistic adversarial loss.

Fréchet inception distance (FID) [24] is the main metric to assess the generation quality. The authors used two sources to estimate the FID; the original images, and quantitative features related to cellular tumors. They decided to include these features since the generated samples should contain meaningful information for medical analysis. Moreover, the experiments related to the latent space showed how the density in the space shifts smoothly according to the amount of cancer cells in the image, and linear interpolations produced consistent tissue transformations.

### Systematic review

This review was carried out following the guidelines of the Preferred Reporting Items for Systematic Reviews and Meta-Analyses (PRISMA 2020) [25] as close as possible (S3 Table). Even though PRISMA 2020 is mainly designed to structure systematic reviews related to medicine, the results of this review could lead to ideas in the context of DL that could benefit the health field. Additionally, we believe that implementing these guidelines enhances the process and outcomes of a paper review independent of its context. This systematic review is not registered in PROSPERO [26] and neither has nor follows any review protocol.

### Eligibility criteria and search strategy

The only source of information selected for the review are papers written in English whose main topic was the implementation or design of GANs. Additionally, the purpose of the GAN must be image augmentation of cell microscopy. Even though I2I translation could be considered an image augmentation method, we decided to define image augmentation as transforming random noise into a generated sample (as in the traditional GAN definition). Papers are legible as long as they perform image augmentation independently of whether it was not the study goal, i.e., an image classification study that used a GAN architecture to augment the dataset is legible and therefore selected for this review.

The selected databases are IEEE Xplore, ScienceDirect, PubMed, arXiv, and bioRxiv. We retrieved studies without limit in the year of publication, meaning that we will consider any publication from the introduction of GANs in 2014. Additionally, we did not include publications cited in the retrieved papers or publications from different sources, considering that our search criteria include the year of publications of GAN. Table 2 summarizes the query search and other search parameters used in each database.

**Table 2.**
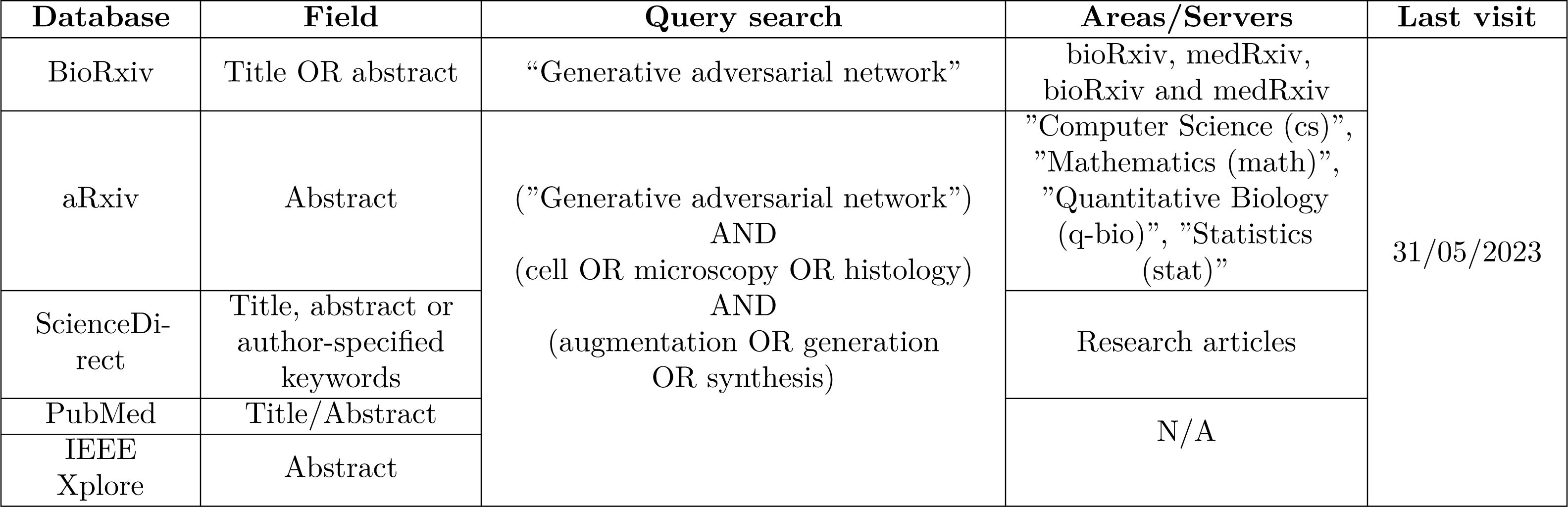
Advanced search query parameters used as search strategy.

A single person was responsible for the whole paper selection process. After removing duplicates, we examined the title and abstract to know if the publication met the eligibility criteria; in case of doubts, the methods and results sections were checked. Papers were considered equally independent of their performance assessment (quantitative, qualitative, specific metric) as long as the publication fulfills the eligibility criteria. Finally, we did not implement any risk of bias tool to the selected studies due to the nature of the field. However, we implemented the ROBIS tool [27] to assess the risk of bias of this systematic review.

### Synthesis methods

Considering that we aim to verify the main applications of GANs in cellular imaging for image augmentation, we decided to use a table as the dominant strategy to summarize the data. The table features are publication, GAN loss, generator (architecture), discriminator (architecture), Performance Metrics, scores, model base-lines, data type, dataset, training type, ablation study, code (availability), designed/modified (D/M), image augmentation as the main task, and notes. Additionally, we designed bar plots when possible to facilitate data visualization.

Depending on the table feature, we considered different conventions to facilitate the data analysis. In the case of data type, we use optical microscopy to represent all datasets acquired with all kinds of light microscopy techniques, including histology or wide-field microscopy. Similarly, the fluorescence microscopy tag includes immunofluorescence images. GAN loss, generator, and discriminator features are simplified using the most representative definitions to minimize the number of possible classes. We decided to keep the performance metrics and scores original, since this study aims to identify the most popular metrics employed in cell imaging. Code, D/M, and image augmentation as main tasks features are binary classes. The first one indicates if the publication has a URL linking to the original implementation, the second one tells if authors created, modified, or redesigned the model architecture or losses from older publications, and the last one tells if the publication goal was to perform image augmentations or if it was an intermediate process. To further synthesize the results, we grouped the studies based on its adversarial loss and briefly discuss their applications, implementations, and common things.

We summarized the publicly available datasets used in the studies that met the legibility criteria similarly to the table described before. The features extracted from each dataset are microscopy modality, number of samples, image resolution, the task for which the dataset was designed for, other published applications of the dataset, binary feature to tell whether the dataset is annotated, cell type, URL, and notes. Moreover, we included a detailed description of each dataset.

## Results

Our methodology retrieved a total of 321 publications and 32 met the eligibility criteria excluding duplicates; 10 (31.25%) from arXiv, 10 (31.25%) from IEEE Xplore, 6 (18.75%) from PubMed, 4 (12.5%) from ScienceDirect, and 2 (6.25%) from bioRxiv. One publication from PubMed had to be excluded since it was inaccessible. Detailed information on the selection process is described in Fig 1.

**Fig 1.**
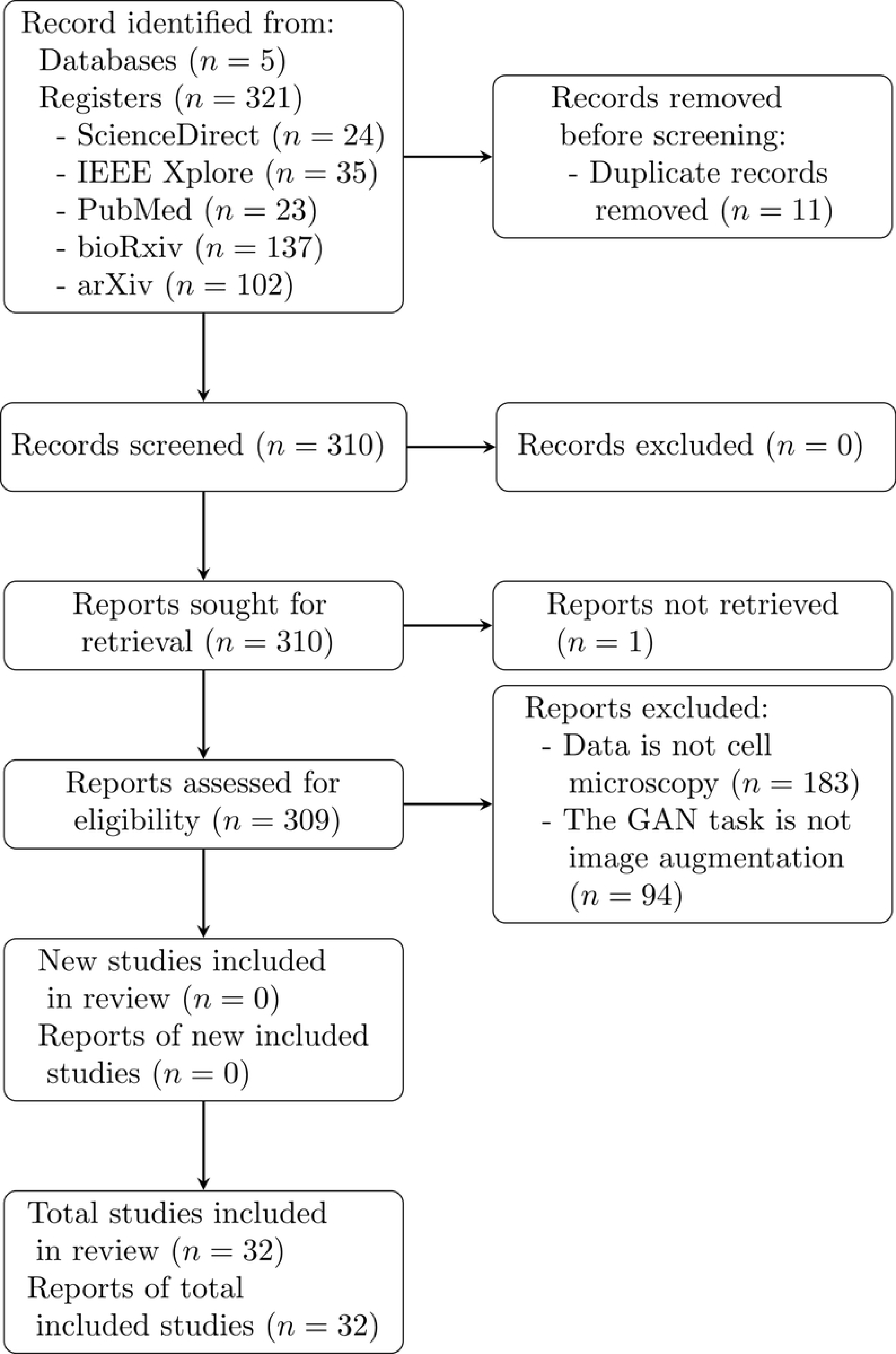
PRISMA search strategy flow diagram.

Considering that we extracted several features from the studies, we decided to analyze the studies from different perspectives using individual features as point of views. Fig 2 visually summarizes the distribution of the studies based on the features we will focus on.

**Fig 2.**
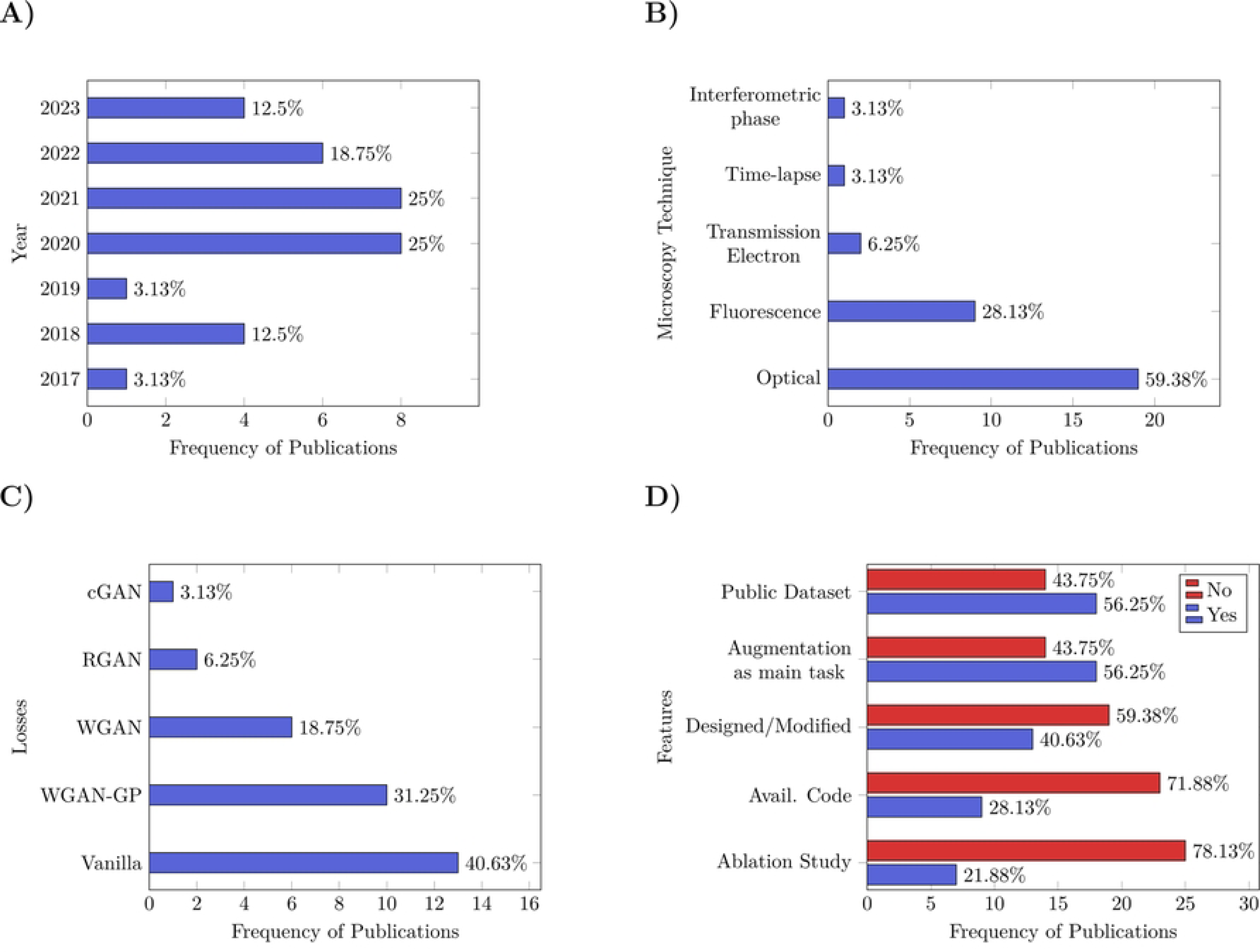
Histograms from the selected studies based on the features used as point of view. **(A)** Yearly publication distribution of GANs used for cell microscopy image augmentation. **(B)** Distribution of microscopy modalities from the datasets used in the selected publications. **(C)** Distribution of adversarial losses. **D)** Distribution of reproducibility and other features.

### Year of publication

We observed an increased interest in data augmentation from 2020, i.e., the year with the most publications (8). Besides 2021 (same amount as in 2020), the second most prolific year is 2022 (6), and then followed by 2023 and 2018, with 4 publications (Fig 2A). The success in 2018 could be attributed to the gained interest in the field due to the introduction of Pix2Pix and CycleGAN in 2017. The oldest publication on cell image augmentation is from 2017. Considering that only a single publication used the Vanilla GAN architectures [28], and that Deep Convolutional GAN (DCGAN) [29] was published in 2016, we suppose that GAN was not powerful enough to produce realistic microscopy images before 2016.

### Microscopy modalities

Fig 2B displays the microscopy modalities of the employed datasets. From there, it is possible to see the dominance of optical and fluorescence microscopy, present in 19 and 9 studies, respectively. There are other three modalities whose only sum up 4 studies, suggesting possible data acquisition limitations for different reasons. Although there is a clear interest in the two microscopy modalities mentioned before, we would like to highlight that the dataset feature is one of the most variables among the classes; the majority of the selected studies use a unique dataset, impeding the comparison between them.

### Adversarial losses

The GAN loss feature has 13 different classes throughout the publications. It is possible to classify losses into two big groups, adversarial and auxiliary. Such losses pretend to guide the architecture training towards the specific task and boost the general performance, respectively. Independent of the data domain and the computer vision task, most GANs use both adversarial and auxiliary losses in their implementation. In this case, however, only 11 publications presented architectures that rely on both types of losses.

The most frequent auxiliary losses are the pixel-wise losses, *L*2 (4 publications) and *L*1 (2 publication) norm. Concerning the adversarial losses, Vanilla GAN loss was the most popular present in 13 publications, followed by the WGAN variants. Fig 2C depicts the adversarial losses’ distribution in the selected studies.

### Vanilla loss

The vanilla adversarial loss, defined by the cross-entropy loss, is implemented so that *G* aims to maximize it; while *D* tries to minimize it. Interestingly, this loss approximates a function dependent on the Jensen Shannon Divergence (*D_JS_*) when *D* reaches its theoretical optimum.

Different from Kullback-Leibler divergence (*D_KL_*), a popular metric for distribution comparison, *D_JS_* is a symmetric statistical distance, and its output bounds to [0, 1], an essential factor to stabilize neural networks training. However, similar to *D_KL_*, *D_JS_* cannot measure the distance between two distributions with disjoint supports. If, for instance, the real and generated distribution do not overlap, *D_JS_* = 1 independently of the distance between them, leading to a possible gradient vanishing during training.

From the 13 studies using vanilla adversarial loss, only three publications used auxiliary losses, and another four used GAN augmentation as an intermediate state to boost the performance of a classification network; the rest focused their work on the augmentation process.

Most studies used traditional architectures such as CNN or vanilla generators and discriminators. Only one used alternative architectures; StyleGAN generator and U-net discriminator [31]. In that work, the authors designed this GAN to generate time series images of two different microscopy modalities simultaneously using a two-branched generator. Another publication presented a two-branched generator to produce images and segmentation masks in a single run with a discriminator that takes a 4-channel input [32].

Kastaniotis et al. used attention maps from a trained classifier to perform knowledge transfer in the discriminator and focus the generator on details [33]. Some publications presented pipelines that combine several GAN architectures [34] [35] [36] [37] and a study that aims to compare the performance of different GAN architectures [38].

### Wasserstein adversarial loss

Given that conventional adversarial losses suffer from training instability by its intrinsic definition, the authors of Wasserstein GAN designed an adversarial loss robust against this instability. As its name implies, WGAN exploits the Wasserstein-1 distance as a loss function. The Wasserstein-1 distance can be elucidated as the minimum amount of work required to move an input distribution to match a target distribution. The main benefit of this measure is that its output is proportional to the distance between the distributions, independently of whether there are disjoint supports. However, its computational complexity makes its calculation infeasible with high-dimensional data.

Because of this, Arjovsky et al. [16] proposed to use the Kantorovich-Rubinstein duality and reformulate the Wasserstein-1 distance into a maximization problem of K-Lipschitz functions [39]. The main challenge of this reformulation is to encourage the neural network architecture to represent K-Lipschitz functions during training. The first approach was to clip the network weights after each gradient update to lie in a compact space. Moreover, WGAN implementation replaces the discriminator with a critic *C* network that outputs a measure instead of a probability. Nevertheless, weight clipping adds a new hyperparameter to the training process and limits the generator generalization power. As a consequence, Gulrajani et al. [40] encourages K-Lipschitz functions with a term addition to the objective function that penalizes the norm of the gradients of *C* instead of limiting the weight values.

Wasserstein loss is the most used adversarial training, with a total of 16 when grouping regular (6) and gradient penalty WGAN (10). Five studies implemented auxiliary losses, eleven used the GAN architectures as intermediate tasks (to boost classification, segmentation, and object detection), and eleven were implementations of already existing models without any modifications. There are studies using PGGAN [41] [42], sinGAN [43], StyleGAN [44], and StyleGAN2 [45] [46]. Bo et al. designed Info-WGANGP, a model using ResNet architectures and an auxiliary network to maximize the mutual information between the images and the generator input to generate realistic images that will later contribute to segmentation and classification architectures [47]; later in 2023, Anaam et al. designed a pipeline based in Info-WGANGP to classify HEp-2 cell images [48]. Osokin et al. trained a WGAN-GP with a Star-shaped generator to train it so that each branch generates a single channel from the fluorescence microscopy target [49]. Finally, Anaam et al. implemented different GAN architectures to evaluate which is the best to do image augmentation and boost the performance of a classification network. They compared DCGAN, WGAN, and WGAN-GP with FID and Classifier two-sample test (C2ST) to generate immunofluorescence images [50].

### Relativistic adversarial loss

The Relativistic GAN (RGAN) is a modification to the GAN losses to measure how much more realistic a real sample is, compared to a generated one. A characteristic of GAN training, is that *D* is always fed with a pool of images where half of them are generated, and the other half are real. Since *D* already knows the amount of real and synthetic data, the probability of classifying correctly a real image should decrease as the generated images start to get classified as real. A benefit of RGAN is that it can be easily applied to those adversarial losses using a regular discriminator. Analogous to WGAN, the discriminator can be seen as a critic with a final sigmoid layer, the critic measures how realistic an input is, while the activation function transforms that value into a probability. The authors used this new critic, subtract the output of real and generated image, and then feed it into the activation layer. In this way, the output of the whole discriminator tells how much more realistic a sample is compared to another one. This implementation has independent objectives for the discriminator and generator that they train to maximize. Moreover, the generator now has an active role in both discriminator and generator objectives. Empirical results suggest that RGAN is more efficient in terms of performance, stability and complexity compared to Vanilla GAN, LSGAN and WGAN [17].

The only two studies using relativistic GAN are a modification of PathologyGAN, a model combining BigGAN, StyleGAN, and RGAN. In [51], the authors included an “inverse generator” to transform images back into the input latent space to extract significant features and generate samples at the same time. On the other hand, the second study implemented an unmodified version of PathologyGAN and selected images based on their latent representation to boost a classification network [52].

### Conditional adversarial loss

Conditional GAN (cGAN) generates images by conditioning its input in several forms, it can use any condition including labels, vectors, or even images. In this way, the generation is controlled after training and researchers produce images with desired characteristics. This review retrieved three using conditional adversarial GANs. The great flexibility of cGAN allows that any adversarial loss can be converted into conditional.

Eschweiler et al. presented a GAN using a 3D U-Net generator with a PatchGAN discriminator to generate 3D fluorescence microscopy images using segmentation masks [53]. Although this work can be considered an I2I translation approach, we decided to include this study in the review because the authors used an algorithm to produce synthetic segmentation masks and generate images using them as generator input.

Kunzmann et al. compared the image quality of a conditional DCGAN against a latent diffusion model to assess the potential of synthetic data [54]. They only used qualitative measures in this study and concluded that latent diffusion models outperform conditional DCGAN.

Dee et al. implemented a conditional StyleGAN2 architecture with the goal of increasing the performance in thyroid histopathology images classification [46]. In another study, Dolezal et al. also trained a conditional StyleGAN2 architecture to generate lung cancer cells and boost a classifier [55]; the objective was to understand the inference process of the classifier through XAI and extract knowledge from the trained network.

### Reproducibility and other features

Only 9 publications shared their implementations, and 18 used public datasets to test their results (some of them also use their own datasets that are not publicly available). On the other hand, 18 publications had image augmentation as the main task of the study, 20 used unmodified GAN architectures published before, only 7 performed an ablation study, and 19 did not use any baseline (Fig 2D). One could argue that since around 60% of the studies used unmodified GANs and 43% of publications focused on a different computer vision task, there is no need to do an ablation study, use a baseline, or share a code implementation. However, we believe that a reference point (baseline and ablation study) is essential to assess the performance of an approach that uses GANs, even for an intermediate task such as data augmentation to boost classification performance.

Regarding network architectures, regular CNN decoder and encoder are the most popular generator and discriminator architectures correspondingly, possibly because a significant part of the studies implemented unmodified GANs and that image augmentation is only an intermediate task. Interestingly, 5 studies used variations of StyleGAN and 3 studies implemented PathologyGAN architecture. Another popular generator architecture was ResNet. Unexpectedly, only 2 publications implemented variations of PatchGAN. The last is surprising to us if we consider that PatchGAN is easily one of the most popular discriminator architectures in computer vision, since it can capture local information that regular discriminators can not. Finally, the majority of architectures used an unsupervised training fashion, with only six supervised and one self-supervised.

There are more than 20 performance metrics, including qualitative assessment and metrics based on tasks different from image augmentation. Some recurrent metrics for image augmentation are Inception score (IS) [30] and FID. The baseline feature is another one with high variance, although a significant part of the studies did not use baselines to compare their results. There are at least 7 different baselines through the pool of publications. Some studies also used baselines towards the principal task to solve, meaning that some baselines are not GAN architectures.

Finally, Table 3 collects the included studies with the most relevant features extracted from the paper. A full version of the table is presented in S1 Table.

**Table 3.**
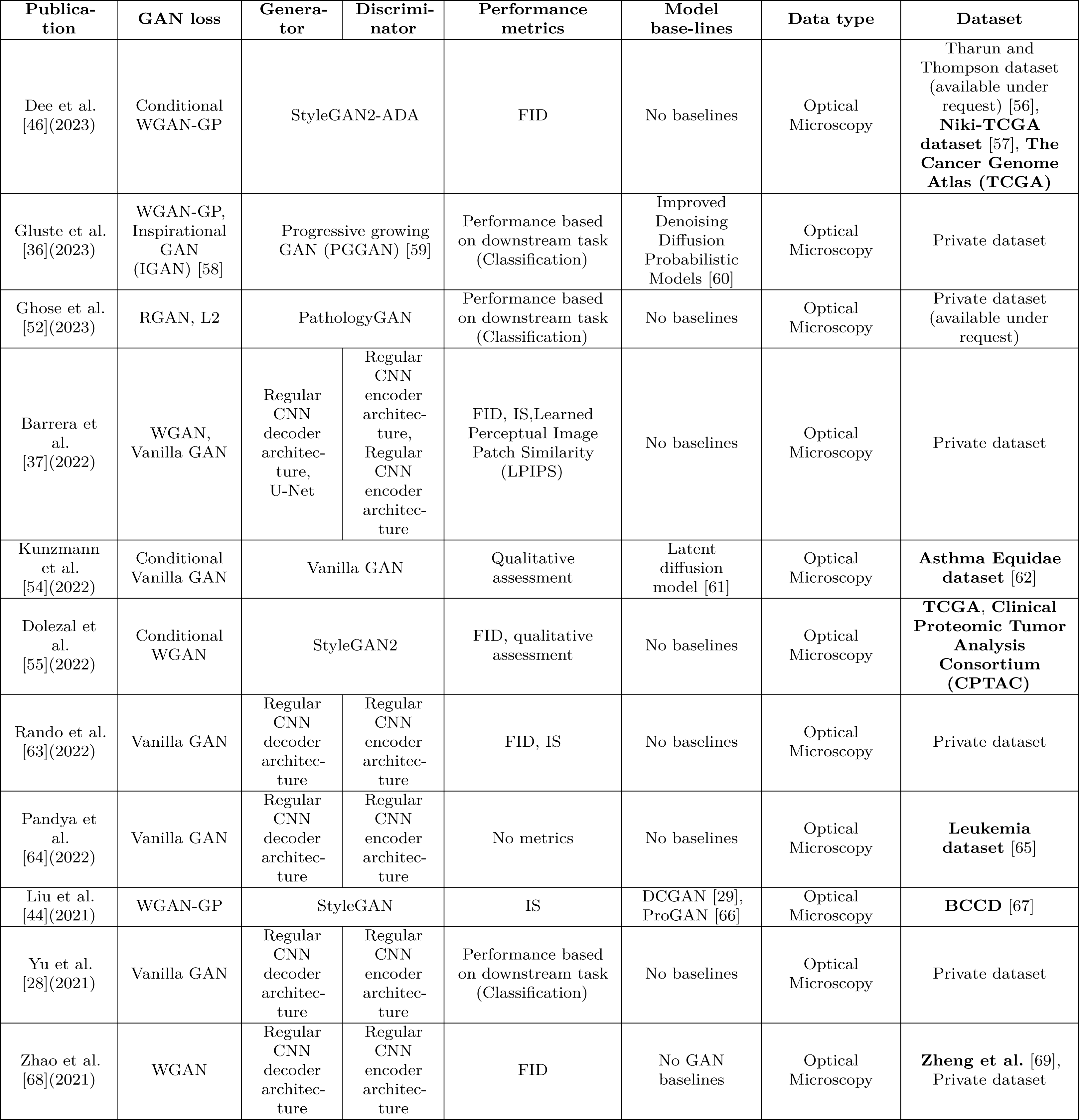
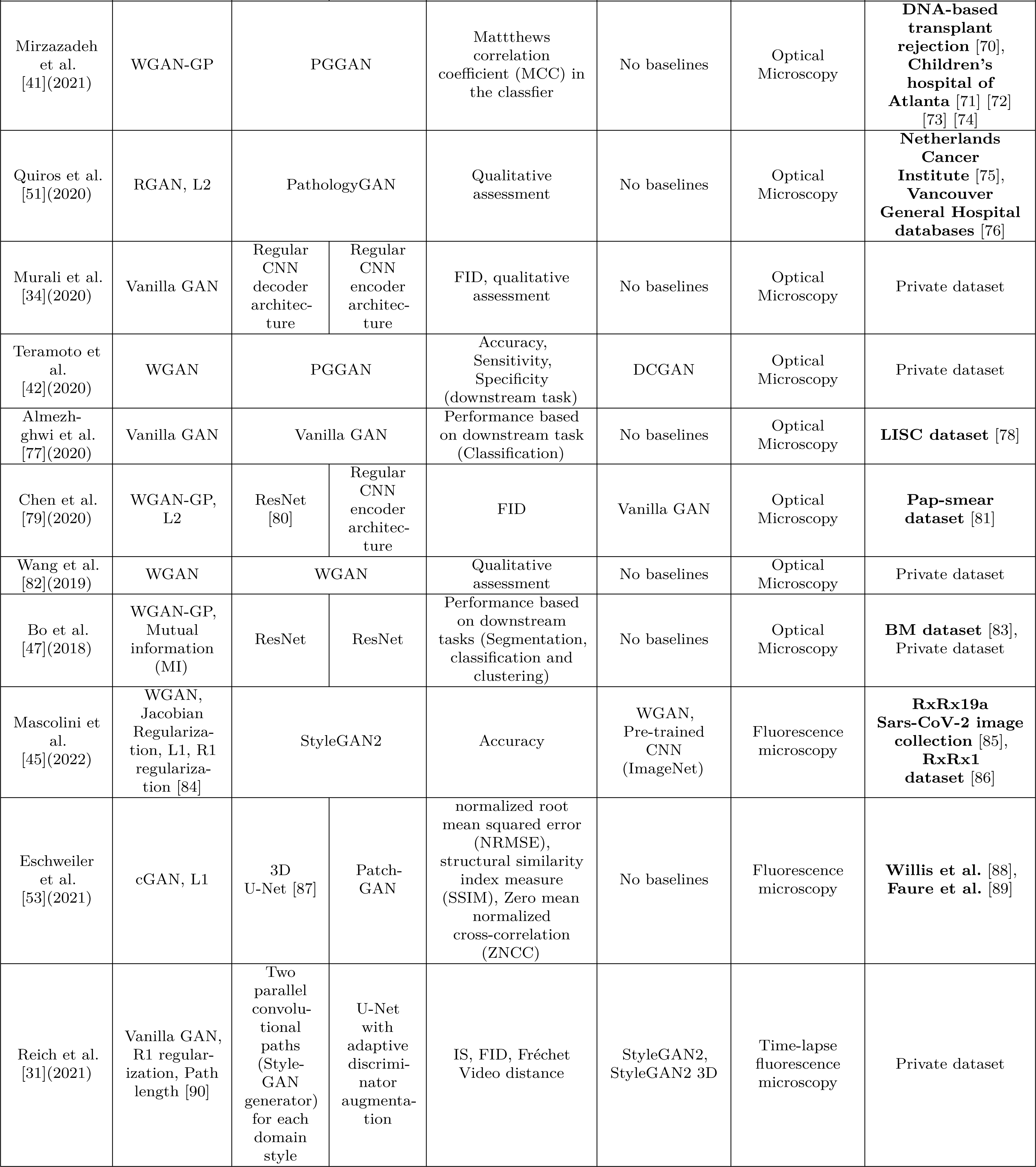
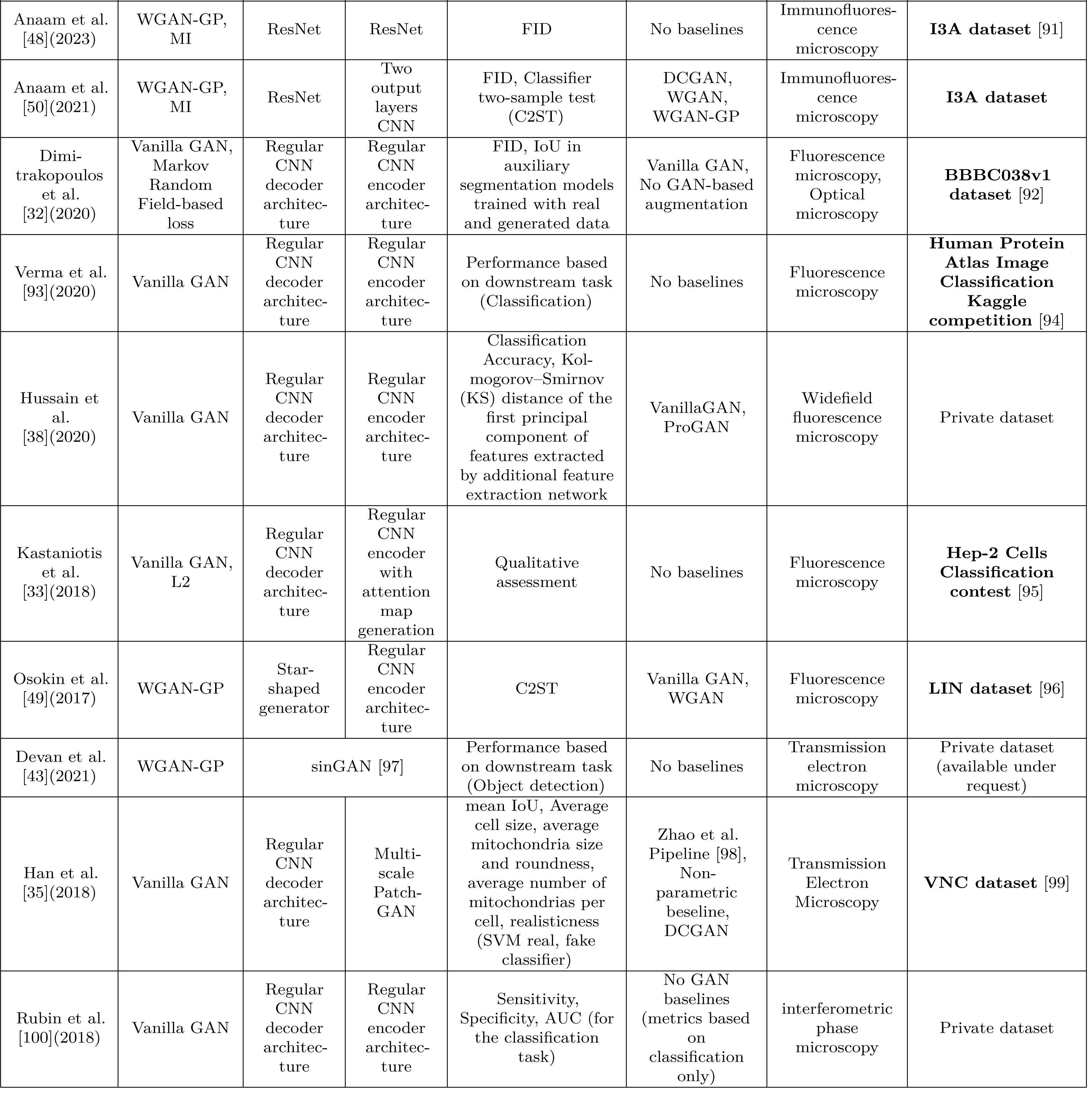
Summary of the studies that met the eligibility criteria. Publicly available datasets are highlighted with bold text.

| Publication | GAN loss | Generator | Discriminator | Performance metrics | Model base-lines | Data type | Dataset |
| --- | --- | --- | --- | --- | --- | --- | --- |
| Dee et al. [46](2023) | Conditional WGAN-GP | StyleGAN2-ADA |  | FID | No baselines | Optical Microscopy | Tharun and Thompson dataset (available under request) [56], **Niki-TCGA dataset** [57], **The Cancer Genome Atlas (TCGA)** |
| Gluste et al. [36](2023) | WGAN-GP, Inspirational GAN (IGAN) [58] | Progressive growing GAN (PGGAN) [59] |  | Performance based on downstream task (Classification) | Improved Denoising Diffusion Probabilistic Models [60] | Optical Microscopy | Private dataset |
| Ghose et al. [52](2023) | RGAN, L2 | PathologyGAN |  | Performance based on downstream task (Classification) | No baselines | Optical Microscopy | Private dataset (available under request) |
| Barrera et al. [37](2022) | WGAN, Vanilla GAN | Regular CNN decoder architecture, U-Net | Regular CNN encoder architecture, Regular CNN encoder architecture | FID, IS, Learned Perceptual Image Patch Similarity (LPIPS) | No baselines | Optical Microscopy | Private dataset |
| Kunzmann et al. [54](2022) | Conditional Vanilla GAN | Vanilla GAN |  | Qualitative assessment | Latent diffusion model [61] | Optical Microscopy | **Asthma Equidae dataset** [62] |
| Dolezal et al. [55](2022) | Conditional WGAN | StyleGAN2 |  | FID, qualitative assessment | No baselines | Optical Microscopy | **TCGA, Clinical Proteomic Tumor Analysis Consortium (CPTAC)** |
| Rando et al. [63](2022) | Vanilla GAN | Regular CNN decoder architecture | Regular CNN encoder architecture | FID, IS | No baselines | Optical Microscopy | Private dataset |
| Pandya et al. [64](2022) | Vanilla GAN | Regular CNN decoder architecture | Regular CNN encoder architecture | No metrics | No baselines | Optical Microscopy | **Leukemia dataset** [65] |
| Liu et al. [44](2021) | WGAN-GP | StyleGAN |  | IS | DCGAN [29], ProGAN [66] | Optical Microscopy | **BCCD** [67] |
| Yu et al. [28](2021) | Vanilla GAN | Regular CNN decoder architecture | Regular CNN encoder architecture | Performance based on downstream task (Classification) | No baselines | Optical Microscopy | Private dataset |
| Zhao et al. [68](2021) | WGAN | Regular CNN decoder architecture | Regular CNN encoder architecture | FID | No GAN baselines | Optical Microscopy | **Zheng et al.** [69], Private dataset |
| Mirzazadeh et al. [41](2021) | WGAN-GP | PGGAN |  | Matthews correlation coefficient (MCC) in the classifier | No baselines | Optical Microscopy | **DNA-based transplant rejection** [70], **Children's hospital of Atlanta** [71] [72] [73] [74] |
| Quiros et al. [51](2020) | RGAN, L2 | PathologyGAN |  | Qualitative assessment | No baselines | Optical Microscopy | **Netherlands Cancer Institute** [75], **Vancouver General Hospital** databases [76] |
| Murali et al. [34](2020) | Vanilla GAN | Regular CNN decoder architecture | Regular CNN encoder architecture | FID, qualitative assessment | No baselines | Optical Microscopy | Private dataset |
| Teramoto et al. [42](2020) | WGAN | PGGAN |  | Accuracy, Sensitivity, Specificity (downstream task) | DCGAN | Optical Microscopy | Private dataset |
| Almezhghwi et al. [77](2020) | Vanilla GAN | Vanilla GAN |  | Performance based on downstream task (Classification) | No baselines | Optical Microscopy | **LISC dataset** [78] |
| Chen et al. [79](2020) | WGAN-GP, L2 | ResNet [80] | Regular CNN encoder architecture | FID | Vanilla GAN | Optical Microscopy | **Pap-smear dataset** [81] |
| Wang et al. [82](2019) | WGAN | WGAN |  | Qualitative assessment | No baselines | Optical Microscopy | Private dataset |
| Bo et al. [47](2018) | WGAN-GP, Mutual information (MI) | ResNet | ResNet | Performance based on downstream tasks (Segmentation, classification and clustering) | No baselines | Optical Microscopy | **BM dataset** [83], Private dataset |
| Mascolini et al. [45](2022) | WGAN, Jacobian Regularization, L1, R1 regularization [84] | StyleGAN2 |  | Accuracy | WGAN, Pre-trained CNN (ImageNet) | Fluorescence microscopy | **RxRx19a Sars-CoV-2 image collection** [85], **RxRx1 dataset** [86] |
| Eschweiler et al. [53](2021) | cGAN, L1 | 3D U-Net [87] | Patch-GAN | normalized root mean squared error (NRMSE), structural similarity index measure (SSIM), Zero mean normalized cross-correlation (ZNCC) | No baselines | Fluorescence microscopy | **Willis et al.** [88], **Faure et al.** [89] |
| Reich et al. [31](2021) | Vanilla GAN, R1 regularization, Path length [90] | Two parallel convolutional paths (StyleGAN generator) for each domain style | U-Net with adaptive discriminator augmentation | IS, FID, Fréchet Video distance | StyleGAN2, StyleGAN2 3D | Time-lapse fluorescence microscopy | Private dataset |
| Anaam et al. [48](2023) | WGAN-GP, MI | ResNet | ResNet | FID | No baselines | Immunofluorescence microscopy | **I3A dataset** [91] |
| Anaam et al. [50](2021) | WGAN-GP, MI | ResNet | Two output layers CNN | FID, Classifier two-sample test (C2ST) | DCGAN, WGAN, WGAN-GP | Immunofluorescence microscopy | **I3A dataset** |
| Dimi-trakopoulos et al. [32](2020) | Vanilla GAN, Markov Random Field-based loss | Regular CNN decoder architecture | Regular CNN encoder architecture | FID, IoU in auxiliary segmentation models trained with real and generated data | Vanilla GAN, No GAN-based augmentation | Fluorescence microscopy, Optical microscopy | **BBBC038v1 dataset** [92] |
| Verma et al. [93](2020) | Vanilla GAN | Regular CNN decoder architecture | Regular CNN encoder architecture | Performance based on downstream task (Classification) | No baselines | Fluorescence microscopy | **Human Protein Atlas Image Classification Kaggle competition** [94] |
| Hussain et al. [38](2020) | Vanilla GAN | Regular CNN decoder architecture | Regular CNN encoder architecture | Classification Accuracy, Kolmogorov-Smirnov (KS) distance of the first principal component of features extracted by additional feature extraction network | VanillaGAN, ProGAN | Widefield fluorescence microscopy | Private dataset |
| Kastaniotis et al. [33](2018) | Vanilla GAN, L2 | Regular CNN decoder architecture | Regular CNN encoder with attention map generation | Qualitative assessment | No baselines | Fluorescence microscopy | **Hep-2 Cells Classification contest** [95] |
| Osokin et al. [49](2017) | WGAN-GP | Star-shaped generator | Regular CNN encoder architecture | C2ST | Vanilla GAN, WGAN | Fluorescence microscopy | **LIN dataset** [96] |
| Devan et al. [43](2021) | WGAN-GP | sinGAN [97] |  | Performance based on downstream task (Object detection) | No baselines | Transmission electron microscopy | Private dataset (available under request) |
| Han et al. [35](2018) | Vanilla GAN | Regular CNN decoder architecture | Multi-scale Patch-GAN | mean IoU, Average cell size, average mitochondria size and roundness, average number of mitochondrias per cell, realisticness (SVM real, fake classifier) | Zhao et al. Pipeline [98], Non-parametric baseline, DCGAN | Transmission Electron Microscopy | **VNC dataset** [99] |
| Rubin et al. [100](2018) | Vanilla GAN | Regular CNN decoder architecture | Regular CNN encoder architecture | Sensitivity, Specificity, AUC (for the classification task) | No GAN baselines (metrics based on classification only) | interferometric phase microscopy | Private dataset |

### Datasets

Table 4 arranges all the datasets cited by the pool of selected publications. This table gathers relevant metadata regarding the images, keynotes, examples of different applications for the dataset, and the URLs to download when available. An extended version of the Cell microscopy datasets table, including the original and other applications can be found in S2 Table.

**Table 4.**
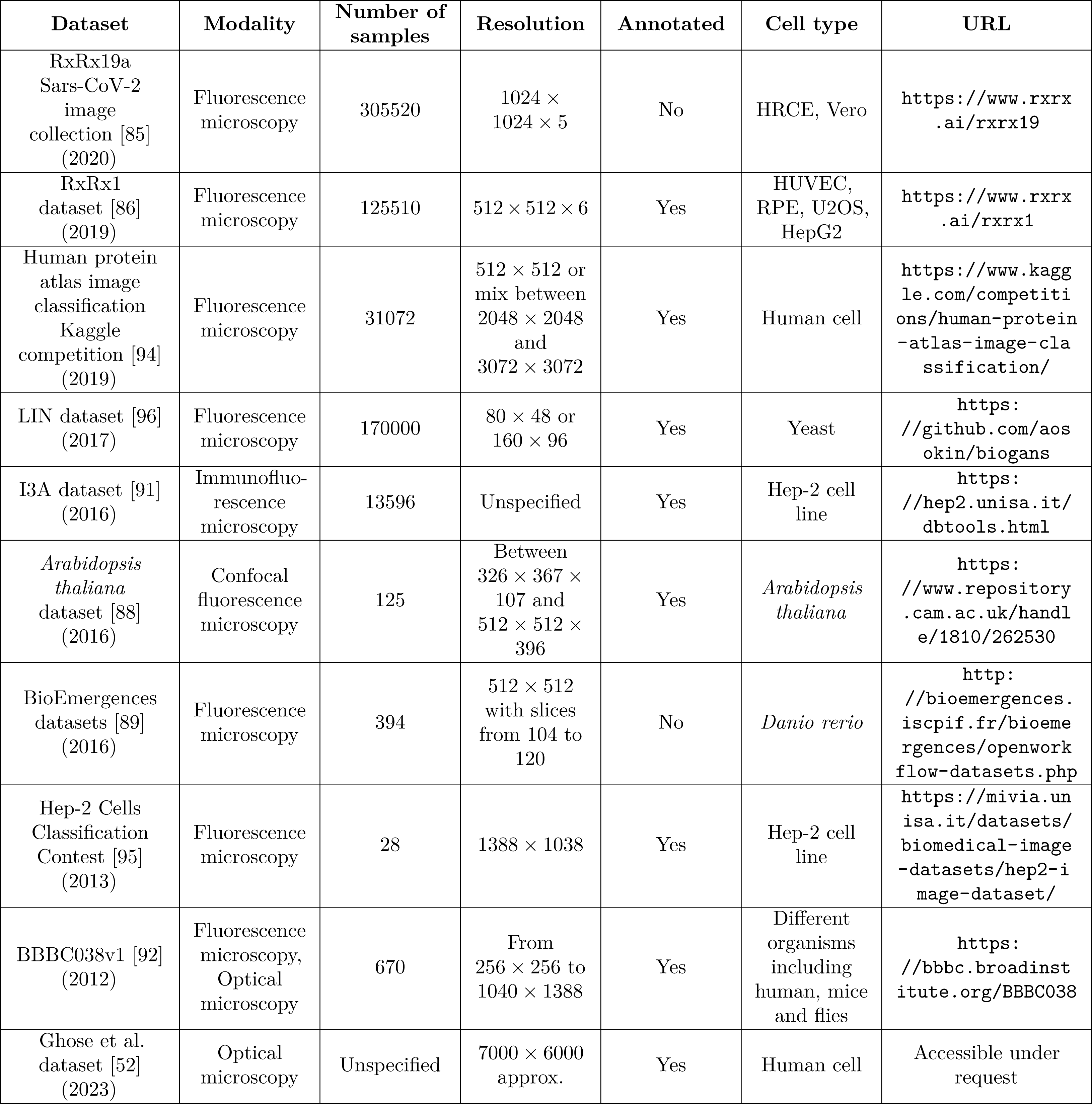
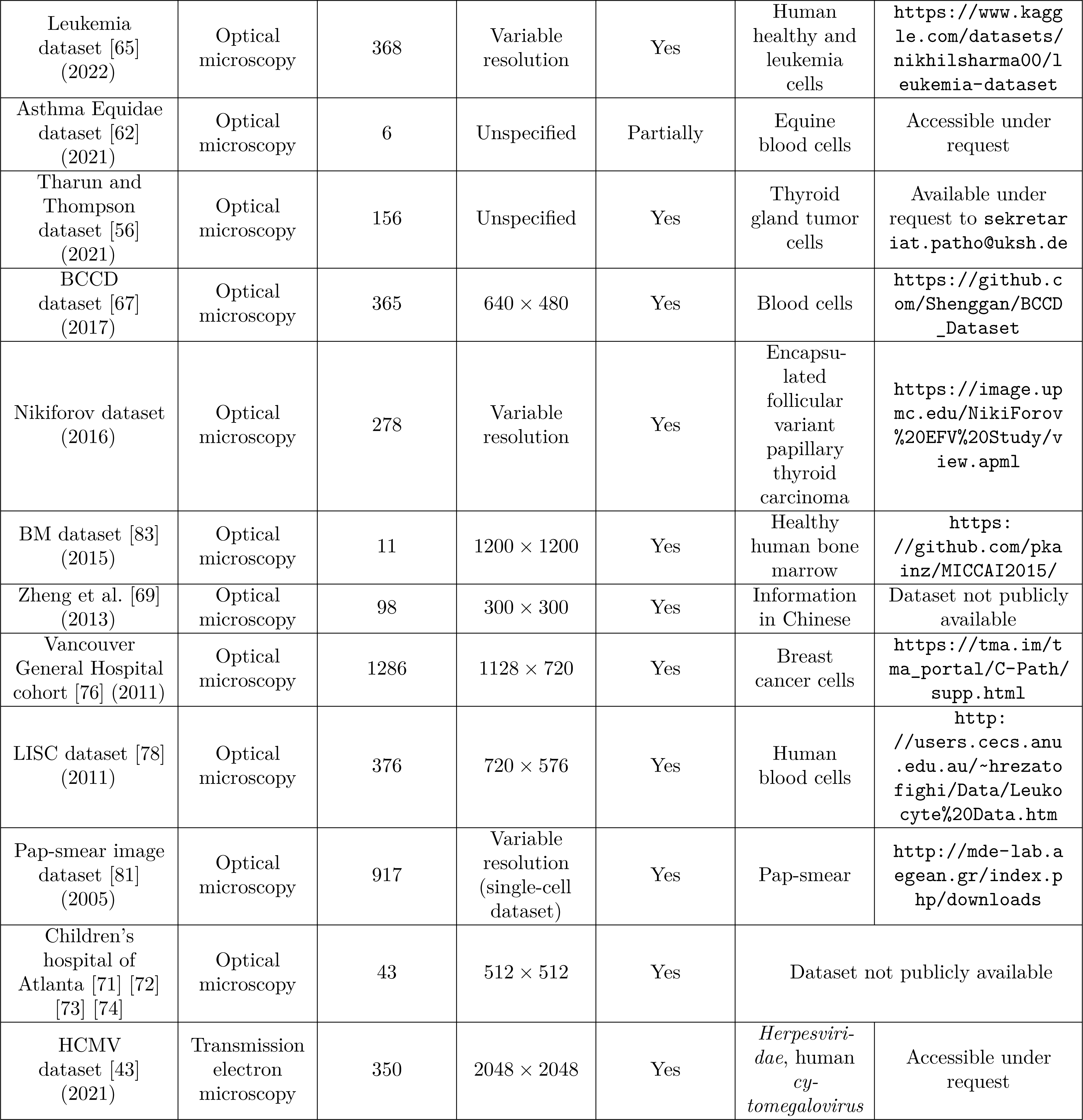
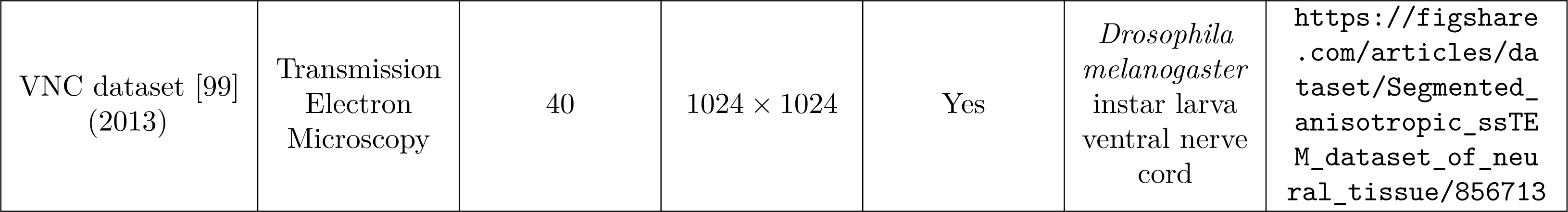
Cell microscopy datasets used in the studies meeting the eligibility criteria.

### RxRx19a

The RxRx19a dataset [85] is the first compilation of cell images to study the morphological effects of COVID-19 in HRCE and Vero cells. It contains a pair of images and a 1024-dimensional vector embedding extracted from a deep-learning approach trained with several cell types and perturbations. It has a total of 305520 fluorescence images of five channels corresponding to different stains. The stains are Hoechst, ConA, Phalloidin, Syto14, and WGA.

### RxRx1

The RxRx1 dataset [86] is a fluorescence microscopy dataset with robust experimental conditions to study biological variations with machine learning. It contains 125510 6-channel images representing 1108 different classes. Each stain highlights different cell organelles; nucleus, endoplasmic reticulum, actin cytoskeleton, nucleolus, mitochondria, and Golgi apparatus. The classes come from gene knockout experiments with small interfering RNA (siRNA), a nucleotide sequence that binds to the messenger RNA (mRNA) and impairs protein synthesis. In the experimental setup of the RxRx1 dataset, a single siRNA targets a single protein, and the 1108 classes correspond to the different siRNAs used by the researchers. Moreover, the experiments were carried out with four different cell types (HUVEC, RPE, U2OS, and HepG2).

### Human protein atlas image classification Kaggle competition

This dataset, composed of 31072 fluorescence microscopy images, is presented by The Human Protein Atlas in a Kaggle competition to build an integrated tool to identify protein locations from high-throughput images. Beyond the expected high classification accuracy, the organizers also included hardware limitations to encourage efficiency and reproducibility.

Users can freely download the dataset on the webpage in two versions, a scaled set of 512 *×* 512 images and a high-resolution version of 2048 *×* 2048 and 3072 *×* 3072 images. This dataset contains 27 cell types with different morphology, and each image is annotated with 28 protein organelle localization labels. From the 4-channel images, contestants were expected to use the green channel to predict the labels, while the other three as references.

### LIN dataset

The LIN dataset compiles 170000 fluorescence images of fission yeast, highlighting two types of proteins. The red channel signals the Bgs4 protein, which localizes the areas of cells’ active growth, while the green channel signals the so-called polarity factors that mark areas of the cells’ cortex. These polarity factors define cell geometry. Each image in the LIN dataset corresponds to a single cell, and the dataset was created to study the interactions between the polarity factors. The dataset targets 41 different polarity factors that control cellular polarity in different ways.

### I3A dataset

This fluorescence dataset has been used in different classification challenges including ICPR 2012, ICIP 2013, and ICPR 2014. It contains HEp-2 cells with a total of 68429 images (13596 training images and 54833 test images). The images were automatically segmented with the DAPI channel and manually annotated by experts. The images are labeled with 6 different classes (homogeneous, centromere, speckled, nucleolar, mitotic spindle, and Golgi) and each sample also has metadata including label, cell intensity, cell mask, and ID. This dataset is only available under request.

### Arabidopsis thaliana dataset

The *A. thaliana* is the product of a publication carried out by Willis et al. [88] where they studied the size and growth regulations of this plant with a Python pipeline. To do so, the authors designed an *A. thaliana* strain with fluorescence markers targeting different genes. There are three channels; the authors used the yellow and red channels to study the cell membranes and nucleus, and the green channel was not analyzed in the study. Each sample corresponds to a stack of volumetric data, which is automatically segmented and partly curated. It has a total of 125 3D stacks (of variable resolution) of six different *A. thaliana* apical meristem.

### BioEmergences datasets

The BioEmergences group is a member of the France-BioImaging National Infrastructure that develops methodologies and tools for the observation, quantification, and modeling of biological processes. In one of their publications, [89], BioEmergences published a workflow to reconstruct cell lineage using 3-dimensional time series fluorescence datasets of stained embryos from Zebra fish (*Danio rerio*), ascidian (*Phallusia mammillata*), and sea urchin (*Paracentrotus lividus*). Each dataset is composed of stain nuclei and membranes 512 *×* 512 stacks with variable 3rd dimension resolution.

Eschweiler et al. [53] only used the Zebra fish data to train their GAN architecture, since they also used the *Arabidopsis thaliana* dataset.

### Hep-2 Cells Classification contest

The MIVIA HEp-2 Images dataset was used in the Cell classification contest carried out in the ICPR 2012. It is an Indirect ImmunoFluorescence (IIF) dataset composed of 28 40*×* images of 1388 *×* 1038 manually labeled and segmented. Each image has metadata including the label (positive, negative, or intermediate intensity), number of objects, number of cells, and number of mitoses.

### BBBC038v1

Kaggle 2018 Data Science Bowl compiled the BBBC038v1 dataset from different laboratories to encourage the development of nuclei segmentation models. It has a total of 841 2D light microscopy images with different staining protocols to increase the variability in the dataset. The whole dataset comprises a total of 37333 segmented nuclei of different cell types and techniques due to the sampling protocol.

### Ghose et al. dataset

This H&E tissue microarray (TMA) dataset is used by Ghose et al. in their image augmentation study [52]. The main characteristic of this dataset is that all samples come from patients with a history of breast cancer event (BCE). The authors extracted the samples from the pathology databases of Oxford University (training cohort) and the Singapore General Hospital (validation cohort). From 133 cases in total, the training cohort (67) comprises one to three TMA cores per patient, while the validation cohort (66) includes three TMA cores per patient. Each H&E TMA core image is approximately 7000 *×* 6000 pixels.

### Leukemia dataset

This blood cell dataset is divided into training and test groups. The training group has 108 images from healthy or ill patients, and each sample is annotated in its file name with a flag to mark the presence of blast cells. Moreover, each image has an associated .xyc file which contains the centroid coordinates of blast cells. The test group has 260 images of single cells, and its file name has a flag to tell whether a blast cell is in the image.

### Asthma Equidate dataset

The samples of this dataset come from Equine bronchoalveolar lavage fluid. The cells were cytocentrifugated and stained with May-Grundwalb Giemsa and then digitalized. Of the six slides, two images are fully annotated, and four are partially annotated. A veterinary pathologist carried out the whole annotation process.

### Tharun and Thompson dataset

This H&E stained dataset is collected from the pathology archives at the University Clinic Schleswig-Holstein and the Woodland Hills Medical Center. Each whole slide image corresponds to a stained section per tumor taken at 40*×* magnification. Each sample was annotated according to the agreement between two pathologists at the whole dataset has five classes: follicular thyroid carcinoma, follicular thyroid adenoma, noninvasive follicular thyroid neoplasm with papillary-like nuclear features, follicular variant papillary thyroid carcinoma, and classical papillary thyroid carcinoma.

### BCCD

The BCCD dataset is a blood cell dataset designed to train cell detection models, it has three labels (red cell, white cell, and platelet) and it is composed of 365 light microscopy 640 *×* 480 images. The images are available in a GitHub repository, where there are data preparation scripts for abnormalities recognition.

### Nikiforov dataset

The samples of this encapsulated follicular variant of papillary thyroid carcinoma (EFVPTC) dataset are taken from 210 patients of different ages diagnosed with noninvasive and invasive EFVPTC. The whole dataset is a collection from 13 different sites in 5 countries. Moreover, each sample was manually curated by 24 thyroid pathologists from 7 countries. The classes in this dataset are: EFVPTC, invasive follicular variant papillary thyroid carcinoma, classical papillary thyroid carcinoma, follicular thyroid carcinoma, and benign.

### BM dataset

The BM dataset is an annotated histology dataset consisting of eleven 1200 *×* 1200 images of human bone marrow. Each image has annotated nuclei, resulting in 4205 detected nuclei. The authors of this dataset developed a cell detection model.

### NIK and VGH cohorts

This H&E-stained histological dataset is a merge of the Netherlands Cancer Institute and Vancouver General Hospital cohorts. The images were samples from 576 patients and, 1286 of them related to breast cancer. The authors of this dataset produced a prognostic model by extracting significant features to assess the prognosis from microscopic image data.

### LISC

The Leukocyte Images for Segmentation and Classification (LISC) dataset is composed of hematological images from healthy subjects. It was thought of as a benchmark for white blood cell classification and nuclei/cytoplasm segmentation techniques. The images come from 8 subjects that produced 100 microscope slides. The experts classified each image with 5 labels (basophil, eosinophil, lymphocyte, monocyte, and neutrophil). Related nucleus and cytoplasm were manually segmented into 250 images.

### Pap-smear image dataset

This dataset has 917 images of human cells stained with the Papanicolau method. Each sample is a single cell with manual annotation inside 7 labels, and it additionally has 20 related numerical features. The authors published this dataset with the idea of a benchmark dataset for classification methods.

### HCMV

The Herpes Human Cytomegalovirus (HCMV) dataset contains 350 transmission electron microscopy (TEM) 2048 *×* 2048 images of human skin fibroblasts infected with HCMV. Each image has bounding boxes annotated by experts. Each box was classified with the three HCMV capsid envelopment stages; naked, budding, and enveloped. Although this dataset is not publicly available, the authors are open to sharing it upon request.

### VNC dataset

The VNC dataset is composed of two stacks of 20 serial section transmission electron microscopy sections of the *Drosophila Melanogaster* third instar larva ventral nerve cord. Only one stack has segmentation annotations highlighting neuron membranes, mitochondria, synapses, and glia/extracellular space.

### Risk of bias with ROBIS

Following the guidelines of ROBIS, we present the concern and rationale for each domains proposed there:

### Concerns regarding specification of study eligibility criteria

#### Low Concern

All signalling questions were answered as “Yes” or “Probably Yes”, so no potential concerns about the specification of eligibility criteria were identified.

### Concerns regarding methods used to identify and/or select studies

#### High Concern

Some eligible studies are likely to be missing from the review, since no additional searching was performed beyond databases. Moreover, only a single person was responsible for screening title, abstract and whole body to classify a study as legible. Finally, only papers written in English were considered.

### Concerns regarding methods used to collect data and appraise studies

#### High Concern

Some bias may have been introduced since only a single person was responsible for data collection, and the nature of the studies do not allow a suitable risk of bias assessment.

### Concerns regarding the synthesis and findings

#### High Concern

The synthesis is likely to produce biased results because it was not possible to consider bias between studies, and the nature of the studies do not allow accounting for variation between them.

#### Risk of bias in the systematic review: High

The main source of risk of bias is that a single person was responsible for the whole review process. However, there is another source of bias and it is the nature of the studies. To our knowledge, there are no available tools or methodologies to assess the risk of bias in DL studies, and not all the ROBIS criteria fit in a systematic review in this field.

## Discussion

### Model implementation

As shown in the results, more than half of the studies used an existing architecture without any modification. In theory, this should not be a problem because the goal of GANs is to learn the probability distribution of the training dataset. However, researchers must consider the nature of the image domain and the requirements of the final goal for those generated images. The most popular applications of medical imaging and cell imaging are diagnosis, evolution assessment, prognosis, or prevention; all of these tasks require high fidelity due to their ethical concerns and possible effects in case of wrong decisions.

One could argue that designing a robust loss objective can be an option to condition the generator and achieve the requirements of this domain. But again, the collected studies indicated that Vanilla GAN is the most popular loss regardless of its theoretical limitations. It could be possible that more sophisticated losses like WGAN could require more resources, but this is not clear from the evidence here. Moreover, most of the auxiliary losses are pixel-wise losses, losses that focus on the pixel level entirely, excluding neighborhood, shapes, textures, and more content features that are essential when the purpose of the architecture is to tell whether a sample looks realistic or not. There are perceptual losses that aim to overcome such limitations [101], but they are more popular for tasks such as I2I translation since it is a supervised loss. It would be beneficial to assess the need for losses capable of capturing features that regular pixel-wise losses can not for these applications.

Additionally, it was surprising to us to see that PatchGAN was present in only two studies. In most cases, a single cell represents a small percentage of the whole image; it is likely that a discriminator evaluates the image as a whole, considering only global features like cell distribution over the medium or cell-to-cell interaction while skipping all local features that could be important, such as cell anatomy. Nevertheless, it is still hard to tell whether PatchGAN or a different architecture has the power to encourage the generator to reproduce those key features, since we evidenced a lack of consensus in the experimental designs.

### Challenges, limitations, and opportunities

#### Experimental design

##### The experimental designing is one of the biggest flaws that we encountered in the legible studies

It is not possible to reproduce, compare, and assess the models presented in the selected studies. Each model is trained and tested with different datasets, each study employs different baselines with distinct performance metrics, and a significant part do not share a code implementation. Even though data augmentation is generally not the main task of the studies, we believe that relying solely on downstream tasks like classification or object detection to assess generative performance should be considered carefully. Such downstream tasks are likely to be performed by other ANNs which are susceptible to bias during training. Supporting the image augmentation evaluation with additional measurements will give a better insight of the image quality, and therefore, ensure adequate performance of downstream architectures.

#### Performance metrics, evaluation, and realism

##### Evaluate and comparing GAN studies is currently not feasible

Another limitation we perceived in this study is how to evaluate GANs performance. One of the reasons for this difficulty is the performance metrics; this study shows how there are different metrics, and although some can be transformed into others, the high number of options limits the assessment between models. Additionally, measuring realism is an intrinsic challenge of generative modeling. Tasks like I2I translation and image augmentation heavily rely on producing realistic images. If it is not possible to accurately measure the generated data quality, it will be unfeasible to compare models or estimate their generalization.

##### Medical and microscopy imaging require more robust metrics than other fields

Some of the most popular metrics, such as IS and FID, are based on ANNs trained with a general-purpose dataset, ImageNet [102], raising the concern of whether such metrics can capture and evaluate the features describing the quality of images that are likely to be significantly different from daily live objects. Considering this concern, Tronchin et al. [103] assess and propose a framework for GANs in the context of medical imaging. We believe that it is essential to focus the attention on how results are measured. A good starting point could be to use the work of Borji [104] as a reference, where he presents and evaluates different qualitative and quantitative performance metrics for GANs.

Even though GANs have been state-of-the-art for image generation for a considerable amount of time, they have important limitations compared to other generative models. First, GANs are known for their training stability; keeping a good balance between the generator and discriminator is mandatory to preserve stable gradients, and they are highly sensible to hyperparameter as well. Second, an intrinsic limitation of these models is mode collapse [105], i.e., when the model cannot learn all the modes of the data distribution and ends up generating samples with limited variability. The current main alternative is Diffusion models [106]. Compared to GANs, Diffusion models are more stable than GANs while better capturing the data distribution at the cost of requiring a considerably higher computational complexity during training and generation. There is already a paper review of Diffusion models in medical imaging [107]. Although it might seem that GANs are losing interest in the community, the adversarial concept is still relevant in computer vision and there is active research on whether merging features of GANs and diffusion models can yield improvements in the field [108].

##### The joint efforts between biology and DL experts is of vital importance

Developing good practices such as including ablation studies, base-lines comparisons, and defining gold standards for metrics and datasets facilitate knowledge transfer. Finally, the risk of bias in this review could be high considering that only a single person was responsible for the searching, screening, and studies analysis, including that we only used 5 databases. However, we must remember that the PRISMA guidelines were not designed for systematic reviews in the field of DL, and some criteria are not applicable. The systematic review is a practice that should be extended to other fields beyond health, but an adaptation is required to make it possible.

## Conclusion

This publication is, to our knowledge, the first systematic review of GANs with cell microscopy for image augmentation. In this study, we examine, summarize, and discuss the most popular methods, together with the current trends in how researchers approach problems in the domain of cell microscopy imaging with GANs. We also compiled and compared the public cell microscopy datasets that have been used for image augmentation since the introduction of GANs to give a brief overview of the available options. Finally, we found some inconsistencies related to the experimental set-up that should be considered not only in studies with GANs, but in any generative modeling study. Reproducibility and comparability is essential to speed up the research progress in any field.

## Supporting information

**S1 Table. Summary of the studies that met the eligibility criteria, full version.**

(PDF)

**S2 Table. Cell microscopy datasets used in the studies meeting the eligibility criteria, full version.**

(PDF)

**S3 Table. PRISMA checklist.**

(PDF)

## Acknowledgments

We thank Miro Mirada (DFKI) for useful discussion and comments on the manuscript.

## Author Contributions

**Conceptualization:** Sheraz Ahmed, Duway Nicolas Lesmes-Leon

**Funding acquisition:** Andreas Dengel

**Investigation:** Duway Nicolas Lesmes-Leon

**Supervision:** Sheraz Ahmed, Andreas Dengel

**Writing – Original Draft Preparation:** Duway Nicolas Lesmes-Leon

**Writing – Review & Editing:** Duway Nicolas Lesmes-Leon, Sheraz Ahmed, Andreas Dengel

## Notes

### Competing Interest Statement

The authors have declared no competing interest.

